# A role for Vps13-mediated lipid transfer at the ER-endosome contact site in ESCRT-mediated sorting

**DOI:** 10.1101/2023.07.06.547997

**Authors:** Sho W. Suzuki, Matthew West, Yichen Zhang, Jenny S. Fan, Rachel T. Roberts, Greg Odorizzi, Scott D. Emr

## Abstract

Endosomes are specialized organelles that function in the secretory and endocytic protein sorting pathways. Endocytosed cell surface receptors and transporters destined for lysosomal degradation are sorted into intralumenal vesicles (ILVs) at endosomes by <u>E</u>ndosomal <u>S</u>orting <u>C</u>omplex <u>R</u>equired for <u>T</u>ransport (ESCRT) proteins. The endosomes (multivesicular bodies, MVBs) then fuse with the lysosome. During endosomal maturation, the number of ILVs increases, but the size of endosomes does not decrease despite consumption of the limiting membrane during ILV formation. Vesicle-mediated trafficking is thought to provide lipids to support MVB biogenesis. However, we have uncovered an unexpected contribution of a large bridge-like lipid transfer protein, Vps13, in this process. Here, we reveal that Vps13-mediated lipid transfer at ER-endosome contact sites is required for the ESCRT pathway. We propose that Vps13 may play a critical role in supplying lipids to the endosome, ensuring continuous ESCRT-mediated sorting during MVB formation.

## INTRODUCTION

Intracellular organelles in eukaryotic cells are essential for compartmentalizing various biochemical and cell signaling reactions. Cells utilize specific vesicle-mediated trafficking systems to populate these organelles with unique sets of protein constituents that carry out these reactions. The ESCRTs (ESCRT-0, -I, -II, -III, and Vps4 AAA-ATPase) sort cell surface receptors and membrane proteins into vesicles that invaginate and bud into the lumen of the late endosome (forming multivesicular bodies, MVBs) (Henne et al., 2011). These MVBs then fuse with the lysosome (the vacuole in yeast) delivering the membrane protein-containing vesicle into the lumen of the lysosome, where they are degraded. In addition to MVB formation, the ESCRTs mediate other critical cellular processes, including the budding of enveloped viruses, such as HIV, cytokinesis, plasma membrane repair, extracellular vesicle formation, and nuclear envelope reformation (Lee et al., 2007; Rusten et al., 2007; Carlton et al., 2008; Hurley., 2015). ESCRT dysfunction has been implicated in numerous diseases, including cancer, neurodegeneration, Huntington’s disease, and Parkinson’s disease (Saksena and Emr, 2009).

Vps13 belongs to a family of lipid transfer proteins that function at various membrane contact sites (Ugur et al., 2020; Dziurdzik and Conibear, 2021; Melia and Reinish, 2022; Neuman et al., 2022). These large proteins (>3,000 AA) have been proposed to bridge membranes to form a direct channel for non-selective lipid transport between two different organelles. The human genome encodes four VPS13 homologs (VPS13A-D), each of which localizes to distinct membrane contact sites. For instance, VPS13A localizes to the ER-mitochondria contact site (Kumar et al., 2018; Yeshaw et al., 2019; Munoz-Braceras et al., 2019), whereas VPS13C localizes to the ER-endosome contact site (Kumar et al., 2018). Mutations in VPS13 homologs have been associated with a variety of neurological disorders, including chorea acanthocytosis (VPS13A) (Rampoldi et al., 2001), Cohen syndrome (VPS13B) (Kolehmainen et al., 2003), Parkinson’s disease (VPS13C) (Lesage et al., 2016), and ataxia (VPS13D) (Seong et al., 2018; Gauthier et al., 2018). Yeast has a single Vps13 that localizes to multiple organelles and membrane contact sites (Dziurdzik and Conibear, 2021). Upon sporulation, Vps13 localizes to the ER-prospore membrane contact site to support spore membrane expansion (Park and Neiman, 2012; Nakamura et al., 2021). Under glucose limitation, it is enriched at the nucleus-vacuole junction (NVJ) (Bean et al., 2018; Lang et al., 2015; Park et al., 2016). It also localizes at the ER-peroxisome and ER-autophagosome contact sites, that function in peroxisome and autophagosome biogenesis, respectively (Yuan et al., 2022; Dabrowski et al., 2023). During cell proliferation, Vps13 primarily localizes to the endosome (Bean et al., 2018; Park et al., 2012; Rzepnikowska et al., 2017). However, the precise function of Vps13 at the endosome remains unclear. Here, we provide evidence that Vps13 forms a lipid transfer channel at the ER-endosome contact site, which is critical for ESCRT-mediated sorting.

## RESULTS

### Vps13 is required for ESCRT-mediated sorting

During endosome maturation, the number of intralumenal vesicles (ILVs) increases. Each endosome contains 60-70 vesicles in yeast. Although the endosomal limiting membrane is consumed to form ILVs, the endosome does not shrink. Vesicle-mediated transport is thought to provide lipids to support MVB biogenesis, but it has not been experimentally tested. To characterize its molecular details, we re-examined the original data from the vacuolar protein sorting (*vps*) mutants, which were isolated because of their defects in the sorting of carboxypeptidase Y (CPY), a soluble vacuolar hydrolase (Robinson et al., 1988). All ESCRT mutants among the original *vps* collection (i.e. *vps4, did4, vps24, vps27, vps20, vps22, vps25, vps28, snf7, vps23,* and *bro1*) exhibited mild sorting defects (Figure S1A). Notably, *vps13* mutants also exhibited a defect similar to that of the ESCRT mutants. Based on this phenotypic similarity, we hypothesized that Vps13 may function in the ESCRT pathway. To test this, we examined the sorting of Mup1, a methionine permease localized to the plasma membrane (PM) (Menant et al., 2006). Upon methionine stimulation, Mup1 is endocytosed and sorted into ILVs at the endosome by ESCRTs (Figure 1A). Then, it is delivered to the vacuole lumen. To evaluate Mup1 sorting, we expressed a GFP-fused Mup1 in yeast cells. In WT cells, Mup1-GFP localized to the PM, but after methionine stimulation, it was sorted into the vacuole lumen (Figure 1B). In contrast, Mup1-GFP was barely sorted in *vps13*Δ cells. Vacuolar delivery of Mup1-GFP results in vacuolar protease-resistant GFP fragments that can be detected by immunoblotting. After 90 min of methionine stimulation, Mup1-GFP was fully processed, and GFP fragments were observed in WT cells, whereas the majority of Mup1-GFP remained as a stable hybrid protein in *vps13*Δ cells (Figure 1C and 1D). We also examined the sorting of carboxypeptidase S (CPS), another cargo of the ESCRT pathway that is delivered from the Golgi and sorted into ILVs (Figure 1E and 1F). Like Mup1-GFP, GFP-CPS was poorly sorted in *vps13*Δ cells. Based on these observations, we conclude that Vps13 is required for ESCRT-mediated sorting.

**Figure 1.**
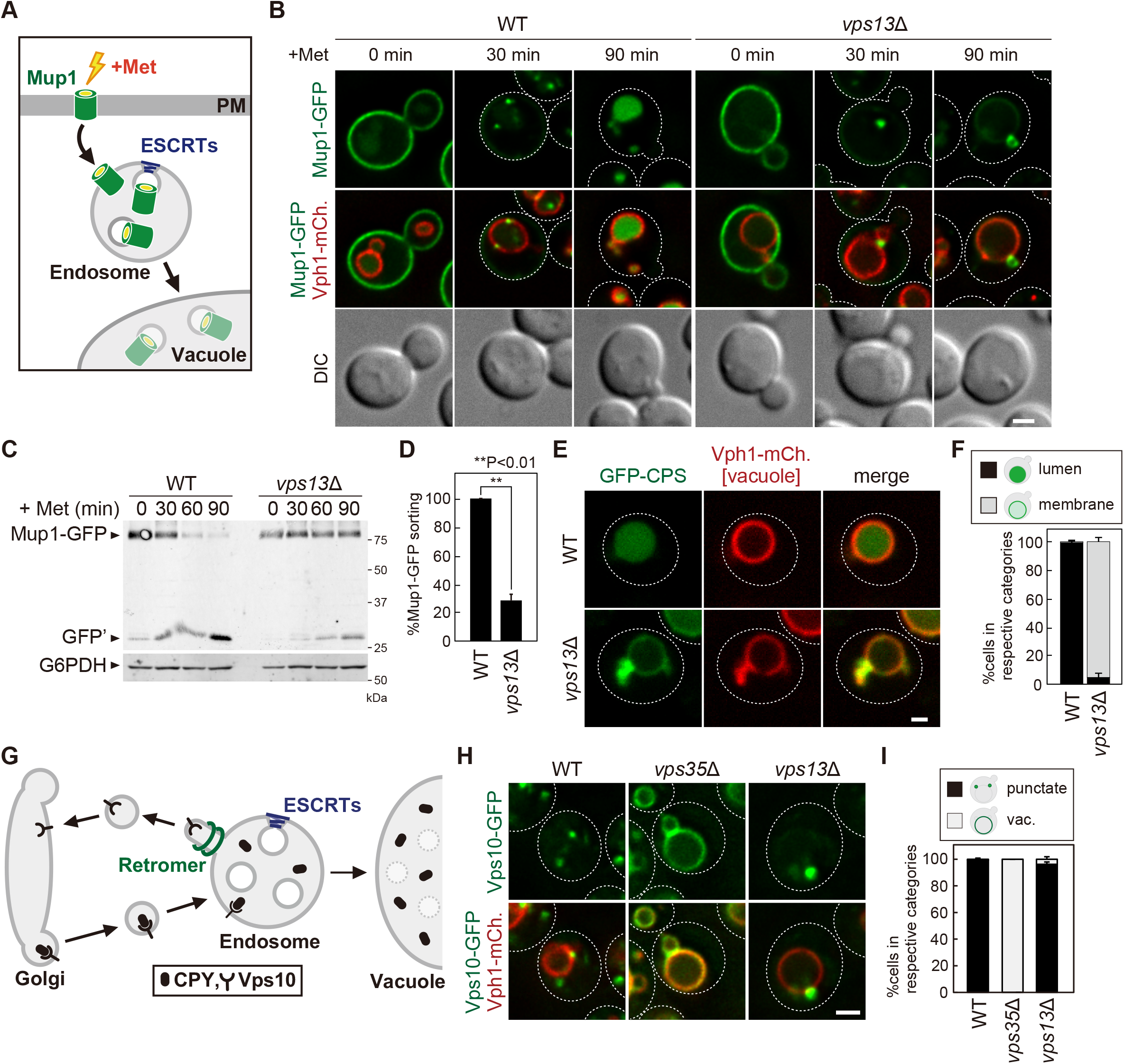
Vps13 is required for the ESCRT pathway. (A) Schematic of Mup1 sorting. (B) Mup1-GFP localization after methionine stimulation. (C) Western blotting analysis of Mup1 sorting. Cell lysates were analyzed by immunoblotting using anti-GFP and anti-G6PDH antibodies. Vacuole delivery of Mup1-GFP yields protease-resistant GFP fragments (GFP’). (D) Quantification of Mup1-GFP processing. (E) GFP-CPS localization. (F) Quantification of GFP-CPS localization of each category. (G) Schematic of retromer-mediated Vps10 recycling. (H) Vps10-GFP localization in WT, *vps35*Δ (retromer), and *vps13*Δ cells. (I) Quantification of Vps10-GFP localization. Scale bar: 1 µm.

### Vps13 is not required for retromer-mediated endosomal recycling

To ask whether Vps13 is specifically required for the ESCRT pathway, we examined its role in endosomal recycling. Vps10 is a transmembrane protein receptor for CPY. After delivery to the endosome, Vps10 is recycled back to the Golgi by the retromer coat complex, which enables Vps10 to carry out another round of CPY sorting at the Golgi (Figure 1G; Marcusson et al., 1994; Seaman et al., 1997; Seaman et al., 1998). Vps10-GFP localized to punctate structures, which were previously reported to be Golgi or endosomes (Figure 1H and 1I; Marcusson et al., 1994). In *vps35*Δ cells, retromer-mediated endosome-to-Golgi retrograde trafficking is impaired, resulting in accumulation of Vps10-GFP at the vacuole membrane. In *vps13*Δ cells, Vps10-GFP was still localized to the punctate structures. We also examined the localization of another retromer cargo, Kex2 (Voos and Stevens, 1998), and its recycling was also not impaired in *vps13*Δ cells (Figure S1B and S1C). These observations suggest that Vps13 is not required for retromer-mediated recycling.

### Vps13-mediated lipid transfer at the ER-endosome contact site is critical for the ESCRT pathway

Vps13 is a lipid transfer protein that localizes to various membrane contact sites. It has a characteristic hydrophobic cavity, which is critical for the lipid transfer reaction (Figure 2A, 2B, and 2C). To define the role of Vps13 in ESCRT-mediated sorting, we examined its localization. Consistent with previous reports (Bean et al., 2018; Park et al., 2012; Rzepnikowska et al., 2017), endogenously expressed Vps13-GFP formed punctate structures, which colocalized with mCherry-Vps21 (Rab5 homolog), confirming its endosomal localization (Figure S2A and S2B). Approximately 10% of Vps13-GFP punctate structures did not colocalize with mCherry-Vps21, presumably because Vps13 also localized to another organelle (i.e., mitochondria, vacuole) as previously reported (Dziurdzik and Conibear, 2021). Vps13 consists of the Extended Chorein, VAB, APT, ATG2_C, and PH domains (Figure 2A; Melia and Reinish, 2022). Since the VAB domain is critical for organelle targeting (Bean et al., 2018), we truncated the C-terminal region including the VAB domain. This mutant (Δ1852-3144) lost endosomal localization (Figure S2C and S2D), indicating that the C-terminal region of Vps13 is required for association with the endosome.

**Figure 2.**
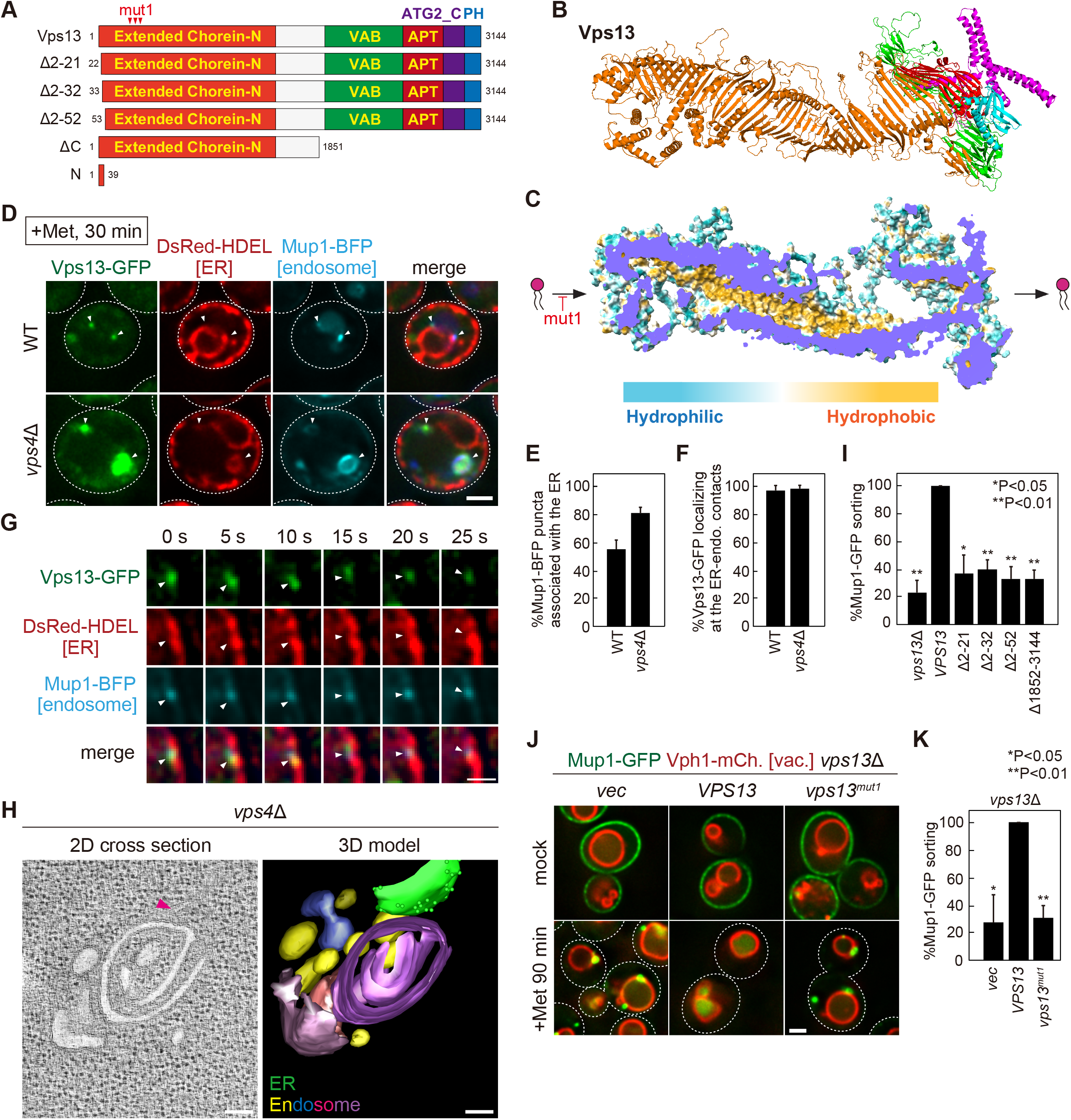
The ER-endosome contact site localization of Vps13 is critical for ESCRT-mediated sorting. (A) Schematic of Vps13 mutants. (B) The ribbon cartoon of Vps13 was generated using AlphaFold2. (C) The surface model of the alphafold predicted structure of Vps13. Blue and yellow indicate hydrophilic and hydrophobic residues, respectively. (D) Vps13-GFP localization at the ER-endosome contact site. Vps13-GFP, DsRed-HDEL (ER), and Mup1-BFP (endosome) expressing WT and *vps4*Δ cells were stimulated with methionine for 30 min. Scale bar: 1 µm. (E) Quantification of Mup1-BFP puncta associated with the ER (DsRed-HDEL). (F) Quantification of Vps13-GFP localization at the ER-endosome contact site. (G) Live-cell imaging analysis of Vps13-GFP at the ER-endosome contact site. Vps13-GFP, DsRed-HDEL (ER), and Mup1-BFP (endosome) expressing WT cells were stimulated with methionine for 30 min. Scale bar: 500 nm. (H) Two-dimensional cross-sections and three-dimensional models of *vps4*Δ cells. ER is traced in green. Ribosomes are indicated as green dots. The endosome stacks are shown in different colors to differentiate individual membranes. Round endosomes are traced in yellow. Larger tubular and cisternal structures are in various shades. Scale bars: 100 nm. (I and K) Quantification of Mup1 sorting in *vps13* mutants. (J) Mup1-GFP localization after methionine stimulation in *vps13* mutants. Scale bar: 1 µm.

Atg2 is also a bridge-like lipid transfer protein that localizes to the ER-autophagosome contact where it provides phospholipids to support autophagosome formation (Osawa et al., 2019; Valverde et al., 2019). The N-terminal 46 residues of Atg2 are sufficient for association with ER membranes and this region can be replaced by the N-terminal 39 residues of Vps13 (Kotani et al., 2018). Therefore, we fused GFP to the N-terminal region of Vps13 (Vps13^1-39^-GFP) and examined its localization. Consistent with previous work (Nakamura et al., 2021), Vps13^1-39^-GFP colocalized with the DsRed-HDEL labeled ER (Figure S2E and S2F), suggesting that the N-terminal region of Vps13 is sufficient for association with the ER membrane.

In humans, VPS13C, a paralog of yeast Vps13, has been shown to localize at ER-endosome contact sites (Kumar et al., 2018). Since the N- and C-terminal regions of yeast Vps13 are required for its association with ER and endosomal membranes, we examined whether yeast Vps13 also localizes to ER-endosome contact sites. For this purpose, we generated yeast cells expressing Vps13-GFP, DsRed-HDEL (ER), and Mup1-BFP. After 30 min methionine stimulation, Mup1-BFP was endocytosed to the endosome (Figure S2G). These Mup1-BFP labeled endosomes were frequently observed in close proximity to the ER, which corresponds to ER-endosome contact sites (Figure 2D, 2E, and S2G). Notably, Vps13-GFP was enriched at these ER-endosome contact sites (Figure 2D, 2F, and S2G). We performed live-cell imaging and revealed that Vps13-GFP puncta stably associated with the ER-endosome contact sites (Figure 2G). We also analyzed the ER-endosome association by live-cell imaging and realized that endosomes dynamically associate and dissociate from the ER membrane (Figure S2H). Therefore, we analyzed Vps13-GFP localization in ESCRT mutants, which exhibit a defect in MVB biogenesis. In ESCRT-defective *vps4*Δ cells, the endosomes were swollen and often observed in close proximity to the ER (Figure 2D and 2E). Vps13-GFP was highly enriched at ER-endosome contact sites in this mutant (Figure 2D and 2F). Electron tomography and three-dimensional modeling of *vps4*Δ cells revealed that the ER membrane was close to the characteristic flattened endosomal structures (known as class E compartments) (Figure 2H and Movie S1). These membranes contacting the endosome were designated ER by the observation of its bound ribosomes, dimensions, and staining by high-pressure freezing and electron tomography (Figure S2I). Notably, ribosomes were excluded from these associated membranes, which was also observed in other contact sites such as the ER-mitochondria contacts (Friedman et al., 2011; Murley et al., 2013). Collectively, these results indicate that Vps13 is localized at ER-endosome contact sites.

To ask if ER-endosome contact site localization is required for ESCRT-mediated sorting, we constructed several Vps13 truncation mutants and examined Mup1 sorting. When we truncated the C-terminal region of Vps13 (Δ1852-3144), which is required for its endosomal localization, Mup1 sorting was impaired (Figure 2I and S2J). Similarly, cells lacking the N-terminal region (Δ2-21, Δ2-32, Δ2-52), required for association with the ER membrane, also exhibited defects in Mup1 sorting. These results suggest that Vps13 association with both the ER and endosome membranes is crucial for the ESCRT pathway.

Vps13 is proposed to bridge two different organelle membranes at a contact site for lipid transport. Consistent with this model, mutations in hydrophobic residues at the Extended Chorein-N domain, responsible for lipid transport, impair sporulation (Figure 2C; Li et al., 2020). We examined Mup1 sorting in this lipid transfer mutant (mut1). It exhibited a severe defect comparable to *vps13*Δ cells (Figure 2J, 2K and S2K), suggesting that the lipid transfer activity of Vps13 is required for ESCRT-mediated sorting.

### Vps13 is required for cargo sorting, but not for cargo ubiquitination and ESCRT recruitment

During ESCRT-mediated sorting, transmembrane cargoes are ubiquitinated and then recognized by ESCRT-0 (Figure S3A; Henne et al., 2012). Subsequently, downstream ESCRTs (ESCRT-I, ESCRT-II, ESCRT-III, and Vps4 AAAase) are recruited to the endosomal surface. ESCRT-III forms a unique spiral structure, that induces membrane invagination and constriction. Finally, Vps4 catalyzes membrane scission. We sought to determine whether Vps13 is required at a specific stage of ESCRT-mediated sorting.

We first examined the ubiquitination status of ESCRT cargoes in *vps13*Δ cells. We immunoprecipitated Mup1-GFP from methionine-stimulated cells and we were able to detect ubiquitinated forms of Mup1-GFP even in *vps13*Δ cells (Figure 3A). To further investigate the requirement of Vps13 in cargo ubiquitination, we used a rapamycin-dependent degradation system (Figure 3B; Zhu et al., 2017). In this system, FKBP-fused cargo proteins and three ubiquitin-conjugated FRBs (FRB-3xUb) are co-expressed. Upon rapamycin treatment, FKBP forms a complex with FRB, which allows ubiquitin recruitment to cargoes. By doing this, it can induce ESCRT-mediated sorting in a ubiquitin ligase-independent manner. After a 90 min treatment with rapamycin, Can1-FKBP was efficiently sorted into the vacuole lumen in WT cells, whereas it was poorly sorted in *vps13*Δ cells (Figure 3C, 3D, and S3B). These results indicate that Vps13 is not required for cargo ubiquitination.

**Figure 3.**
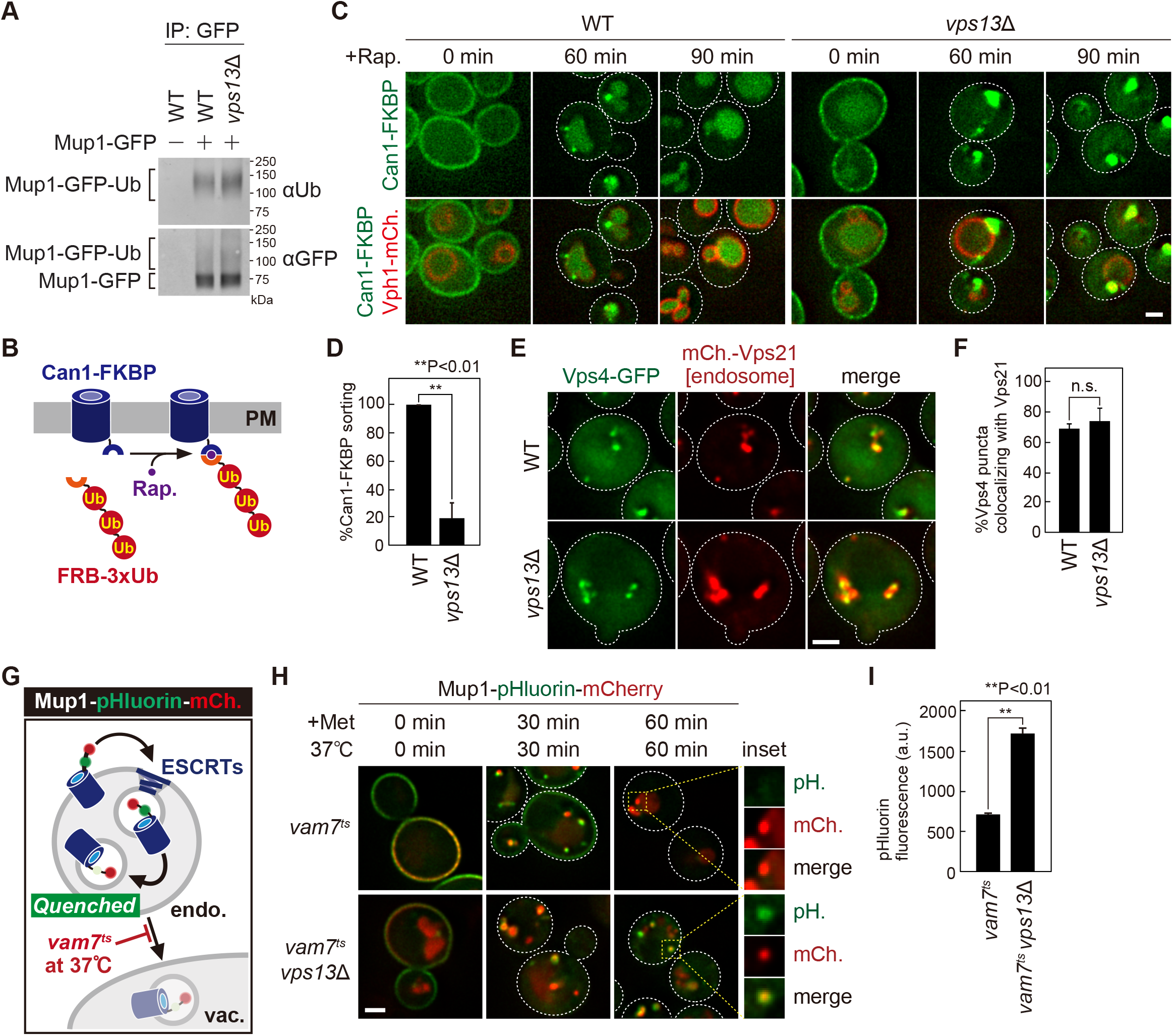
Cargo sorting into the intralumenal vesicle requires Vps13. (A) The ubiquitination of Mup1-GFP. Mup1-GFP expressing cells were stimulated with methionine for 15 min and then immunoprecipitated under denatured conditions. The immunoprecipitated (IP) products were analyzed by immunoblotting using anti-GFP and anti-ubiquitin antibodies. (B) Schematic of a rapamycin-dependent degradation system for Can1. (C) Can1-FKBP localization. Rapamycin-insensitive mutant cells (*tor1-1*, *fpr1*Δ) expressing Can1-GFP-2xFKBP (Can1-FKBP) and FRB-3xUb were treated with rapamycin. (D) Quantification of Can-FKBP sorting. (E) Vps4-GFP localization. (F) Quantification of endosomal Vps4-GFP localization. (G) Schematic of Mup1-pHluorin-mCherry sorting. (H) Mup1-pHluorin-mCherry localization. Mup1-pHluorin-mCherry expressing *vam7^ts^* and *vam7^t^ vps13*Δ cells were incubated at 37°C and stimulated with methionine. (I) pHluorin fluorescence of Mup1-pHluorin-mCherry. Scale bar: 1 µm.

We next examined whether ESCRTs are properly localized to the endosome in *vps13*Δ cells. GFP-Vps27 (ESCRT-0), Snf7-GFP (ESCRT-III), and Vps4-GFP were colocalized with the endosome marker mCherry-Vps21 even in *vps13*Δ cells (Figure 3E, 3F, S3C, and S3D), suggesting that Vps13 function may be downstream of ESCRT recruitment.

Finally, to evaluate cargo sorting into ILVs we constructed a Mup1-pHluorin-mCherry fusion protein (Figure 3G). When this construct is internalized into the ILV, the pHluorin signal is quenched, allowing us to evaluate the status of cargo sorting at the endosome. We expressed this construct in a temperature-sensitive *vam7^ts^*SNARE mutant (Rieder and Emr, 1997; Sato et al., 1998), which has a defect in the fusion of the endosome with the vacuole (Figure 3H and 3I). After 60 min of methionine stimulation, this construct showed punctate structures with mCherry signal but little pHluorin signal, presumably because it was sorted into the ILV. On the other hand, in *vps13*Δ cells, even after 60 min of stimulation, the pHluorin signal of these punctate structures was less quenched. These observations suggest that Vps13 is required for cargo sorting into the ILV, but is not essential for cargo ubiquitination and ESCRT recruitment.

### Vps13 is required for proper ESCRT function at the endosome

To investigate ILV formation in *vps13*Δ cells, we employed electron tomography and three-dimensional modeling (Figure 4A, 4B, and S4A; Movie S2, S3, and S4). Strikingly, *vps13*Δ cells exhibited an increased number of endosomes (MVBs) (Figure 4C and S4B), and these endosomes frequently clustered together (Figure 4A, 4D, S4A, and S4C). Consistent with the electron tomography analysis, fluorescence microscopy also revealed that Vps55-GFP labeled endosomes form clusters in *vps13*Δ cells (Figure 4E and 4F). The size of endosomes in *vps13*Δ cells was 1.78-fold larger than that of WT cells (Figure 4G and S4B), while the number of ILVs per endosome was similar (Figure 4H and S4B). Notably, the size of ILVs in *vps13*Δ cells was 1.34-fold larger and exhibited greater heterogeneity, contrasting with the smaller and more uniform ILVs observed in WT cells (Figure 4I and S4B). Intriguingly, several ILVs in *vps13*Δ cells exceeded 90 nm in size, which was rarely observed in WT cells. Furthermore, we analyzed the inward budding profile (BP) and found that *vps13*Δ exhibited a lower BP frequency compared to WT cells, although the ILV size is larger (Figure 4B, 4J, and S4B). Taken together, these observations suggest that Vps13 is required for appropriate MVB formation.

**Figure 4.**
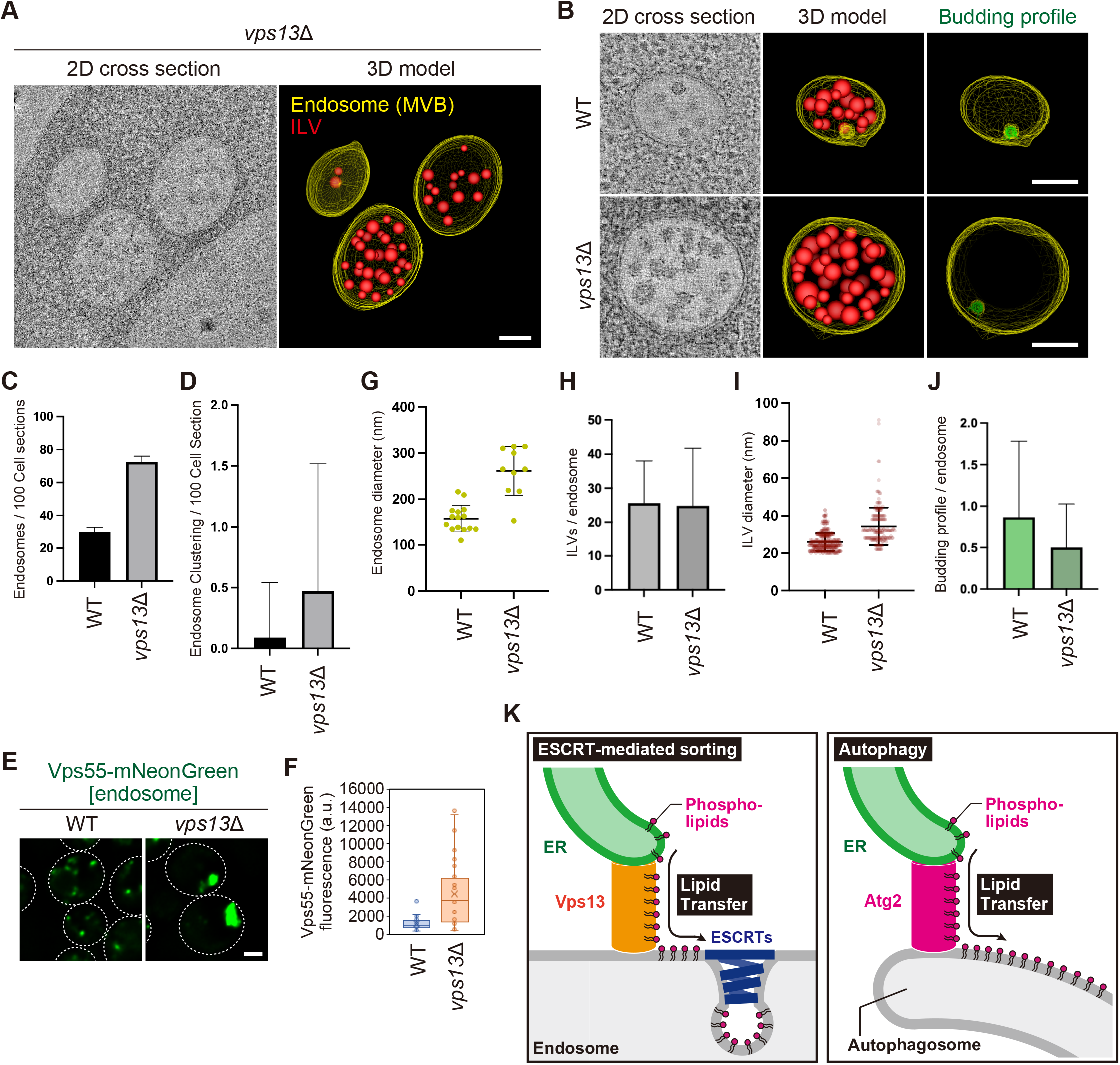
Efficient intralumenal vesicle formation requires Vps13. (A and B) Cross-sectional tomographic slices and three-dimensional models of WT and *vps13*Δ cells. Endosome-limiting membranes are traced in yellow, and detached ILVs are traced in red. ILV budding profiles are traced in green. Scale bars: 100 nm. (C) Quantification of the number of endosomes. (D) Quantification of endosome clustering. (E) Vps55-mNeonGreen localization. Scale bars: 1 um. (F) Fluorescence intensity of the Vps55-mNeonGreen punctate structures. (G) Measurement of the diameter of endosomes. (H) Quantification of the number of ILVs per endosome. (I) Measurement of ILV diameter. (J) Quantification of the budding profile per endosome. (K) Model of a role of Vps13 in ESCRT-mediated sorting. During autophagosome formation, Atg2 provides lipids from ER to the autophagosome to support its expansion.

## DISCUSSION

During MVB formation, the endosome does not shrink, although its limiting membrane is consumed to form ILV. How lipids are supplied to support MVB formation remains an open question. In this study, we found that Vps13 is required for ESCRT-mediated sorting at the endosome. Mutational analysis revealed that Vps13-mediated lipid transfer at the ER-endosome contact site is required for the ESCRT pathway. Cell biological and electron tomography analysis revealed that MVB sorting was impaired in *vps13*Δ cells. Based on these observations, we propose that the limiting membrane of the endosome is consumed during MVB formation, and Vps13 may play a critical role in providing lipids to the endosome that permit continuous ESCRT-mediated sorting (Figure 4K). Strikingly, Atg2, another Vps13-like lipid transfer protein, delivers phospholipids from the ER to the autophagosome to support its expansion (Osawa et al., 2019; Valverde et al., 2019). A similar mechanism may exist in MVB biogenesis.

Vps13 facilitates lipid transport at a much faster rate (34-3400 lipids/sec) than other lipid transfer proteins such as OSBP (<1 lipid/sec) (Zhang et al., 2022). Although this fast lipid flux is ideal for vesicle biogenesis, lipid transfer alone is not sufficient for this process. During autophagosome biogenesis, the lipid scramblase Atg9 plays a critical role in allowing lipids to move across the bilayer from the outer to the inner leaflet (Matoba et al., 2020; Maeda et al., 2020). Interestingly, mammalian VPS13A, which is localized at the ER-PM contact site, interacts with the lipid scramblase XK (Park et al., 2020; Park JS et al., 2022; Guillen-Samander et al., 2022). Vps13 function in the ESCRT pathway may also require a lipid scramblase. Identification of such a lipid scramblase is an important next step in understanding the mechanisms behind MVB biogenesis.

In this study, we propose that lipids supplied by Vps13 are crucial for ESCRT-mediated sorting, but their precise physiological significance remains unclear. Interestingly, electron tomography and three-dimensional analysis revealed that *vps13*Δ cells exhibited a decrease in ILV budding profiles, an increase in ILV size, and a more heterogeneous size distribution. Maintaining appropriate lipid composition might, therefore, impact the efficiency of ILV formation in addition to being important for ILV cargo sorting by the ESCRT machinery.

Vps13 mediates bulk lipid flow between different organelles. Since the lipid composition of these organelles is different, the lipid transfer reaction of Vps13 must be tightly regulated. We demonstrated that Vps13 stably localized to the endosome, but its association with the ER membrane was transient. Consistent with our finding, recent in situ EM analysis revealed that the N-terminus of mammalian VPS13C also appeared to be detached from ER in most cases, while its C-terminus associates with the endolysosomal membrane (Cai et al., 2022). The lipid transfer activity of Vps13 might be regulated by the association with the ER membrane. Understanding how the lipid transfer activity of Vps13 is regulated is an essential issue to be addressed in a future study.

Our study sheds light on a critical role for Vps13 in the ESCRT pathway. Human cells possess ten VPS13-like proteins, including VPS13A/B/C/D, ATG2A/B, Hobbit, Tweek, SHIP164, and UHRF1BP1, all of which function as bridge-like lipid transfer proteins (Toulmay et al., 2022; Levin., 2022). These proteins not only localize to the ER-endosome contact site but also to various other sites, including those between the ER and PM, lysosomes, and autophagosomes. Interestingly, ESCRTs function on diverse membranes, such as the nuclear membrane, PM, lysosomal membrane, and autophagosome. The bridge-like lipid transfer proteins might contribute to ESCRT-mediated membrane remodeling on diverse organelle membranes. Further studies are required to address this fascinating question and to unravel the precise role of these bridge-like lipid transfer proteins in cellular membrane dynamics.

### Limitations of this study

How the membrane is supplied for continuous ILV formation is one of the fundamental questions in cell biology. In this study, we investigated the responsible molecule and identified Vps13 as a potential candidate. However, the contribution of Vps13-mediated lipid transfer to membrane supply cannot be investigated by simply characterizing *vps13*Δ cells, as these cells also exhibit a defect in ILV formation. The development of specific tools to monitor lipid transfer by the Vps13 system is essential to further explore this question.

## ACKNOWLEDGMENT

We thank all Emr lab members for their helpful discussions. We also thank Dr. Martin Graef for sharing plasmids and Dr. Yoshitaka Moriwaki for critical advice for alphafold2 prediction. S.W. Suzuki is supported by Osamu Hayaishi Memorial Scholarship for Study Abroad. This work was supported by a Cornell University Research Grant (CU563704) to S.D. Emr.

## AUTHOR CONTRIBUTIONS

Conceptualization, S.W.S; Methodology, S.W.S; Investigation, S.W.S, M.W., J.S.F., R.T.R., and Y.Z; Writing - Original Draft, S.W.S; Writing - Review & Editing, S.W.S; Funding Acquisition, S.D.E; Resources, S.W.S; Supervision, S.W.S

## MATERIAL AND METHODS

### Yeast Strain and Media

*S. cerevisiae* strains used in this study are listed in Table S1. Standard protocols were used for yeast manipulation (Kaiser et al., 1994). Cells were cultured at 26°C to mid-log phase in YNB medium [0.17% (w/v) yeast nitrogen base w/o amino acids and ammonium sulfate, 0.5% (w/v) ammonium sulfate, and 2% (w/v) glucose] supplemented with the appropriate nutrients.

### Plasmids

Plasmids used in this study are listed in Table S2.

### Antibodies

For immunoblotting, mouse monoclonal anti-GFP (B-2; Santa Cruz, Sc-9996), rabbit polyclonal anti-G6PDH (Sigma-Aldrich, SAB2100871), and anti-ubiquitin (P4D1; Cell Signaling, #3936) were used at dilution factors of 1:5000, 1:10,000 and 1:1000, respectively.

### AlphaFold Structure Prediction

For structure prediction of the full-length of *Saccharomyces cerevisiae* Vps13, we first modeled two segments of Vps13 (1-1860 and 1351-3144 a.a.) by AlphaFold v2.0 and v2.1.1 (Jumper et al., 2021). Then, using the overlapping region (1351-1860 a.a.), we aligned them to generate a full-length of structure.

### Cargo Sorting Assay for the ESCRT pathway

For Mup1 sorting, Cells expressing Mup1-GFP were grown in YNB (-Methionine) media to mid-log phase at 26°C and then treated with 20 µg/ml methionine to stimulate Mup1 sorting. For Can1-FKBP sorting, rapamycin-insensitive mutant cells (*tor1-1*, *fpr1*Δ) expressing Can1-GFP-2xFKBP (Can1-FKBP) and FRB-3xUb were grown to the mid-log phase at 26°C and treated with 200 ng/ml Rapamycin to stimulate its sorting.

### Electron Tomography and Three-dimensional Modeling

Haploid yeast cells were high-pressure frozen and freeze substituted as previously described (Nickerson et al., 2006; Tseng et al., 2021). Liquid cultures were harvested at mid-logarithmic phase, vacuum-filtered on 0.45-μm millipore paper, loaded into 0.5-mm aluminum hats, and high-pressure frozen with a Wohlwend HPF (Wohlwend, Switzerland). Cells were freeze-substituted in an Automated Freeze-Substitution machine (AFS, Leica Vienna, Austria) at −90°C in an en bloc preparation of 0.1% uranyl acetate and 0.25% glutaraldehyde in anhydrous acetone. Samples were then washed in pure anhydrous acetone, embedded in Lowicryl HM20 resin (Polysciences, Warrington, PA), UV polymerized at −60°C warming slowly over 4 days to room temperature. These methods preserve membrane and protein structure and provide consistent en bloc staining (Giddings et al., 2003, Staehelin communications).

A Leica UC6 Ultra-Microtome was used to cut and place serial sections on Formvar-coated rhodium-plated copper slot grids (Electron Microscopy Sciences). 80 nm thin serial sections were cut for transmission electron microscopy (TEM) and 200 nm thick serial sections were cut for dual-axis tomography. Thin sections were imaged with a FEI Tecnai T12 Spirit electron microscope equipped with a 120 kV LaB6 filament and AMT (2 k × 2 k) CCD. TEM of hundreds of cells per strain were used to quality control freezing, embedding, and staining as done previously (Wilson et al., 2021). Thick sections were labeled with fiduciary 15-nm colloidal gold (British Biocell International) on both sides and tilt imaged with a Tecnai 30 (f-30, 300 kV; FEI-Company, Eindhoven, the Netherlands) with dual–tilt series images collected from +60° to −60° with 1.5° increments using a Gatan US4000 4k × 4k charge-coupled device camera (Abingdon, United Kingdom). The tilt series were imaged primarily at 20,000 times magnification and repeated with a 90° rotation to create a dual-axis tomogram with a 3 nm resolution and a 0.4306 nm pixel (Mastronarde., 1997).

Tomograms were built and modeled using the IMOD software package (Kremer et al., 1996, University of Colorado Boulder) using an iMac (Apple). MVB membrane models from dual-axis electron tomograms were manually assigned from the inner leaflet every 5 nm and calculated using IMODmesh. Budding Profiles (BPs) were designated by their negative curvature, since the majority of endosome limiting membrane curvature is positive or spherical in shape. BP models are drawn from the 0° rim at the outer leaflet, measured, and sorted by surface area using only BPs that have more than 750 nm^2^ or approximately half of the mean ILV surface (Wemmer et al., 2011). ILVs are spherical and measured using sphere-fitting models from the vesicle’s outer leaflet (the inner leaflet of the MVB limiting membrane) and ILV diameters were measured using these sphere models. To determine MVB number and MVB clustering, 100 random cell profiles of WT and *vps13*Δ cells were quantified by thin-section electron microscopy. Videos were made using IMOD and QuickTime Pro (Apple). Data were analyzed and graphed using Prism 9 (GraphPad).

### Fluorescence Microscopy

Fluorescence microscopy was performed using a CSU-X spinning-disk confocal microscopy system (Intelligent Imaging Innovations) or a DeltaVision Elite system (GE Healthcare Life system).

A CSU-X spinning-disk confocal microscopy system is equipped with a DMI 6000B microscope (Leica), 100×/1.45 numerical aperture objective, and a QuantEM electron-multiplying charge-coupled device (CCD) camera (Photometrics). Imaging for yeast cells was done at room temperature in YNB medium using GFP and mCherry channels with different exposure times according to each protein’s fluorescence intensity. Images were analyzed and processed with SlideBook 6.0 software (Intelligent Imaging Innovations).

A DeltaVision Elite system is equipped with an Olympus IX-71 inverted microscope, DV Elite complementary metal-oxide semiconductor camera, a 100×/1.4 NA oil objective, and a DV Light SSI 7 Color illumination system with Live Cell Speed Option with DV Elite filter sets. Imaging was done at room temperature in YNB medium using GFP, mCherry, and DAPI (for BFP) channels with different exposure times according to each protein’s fluorescence intensity. Image acquisition and deconvolution (conservative setting; five cycles) were performed using DeltaVision software softWoRx 6.5.2 (Applied Precision)

### Immunoprecipitation under denature conditions for yeast cell lysate

To analyze the ubiquitination status of Mup1, cells expressing Mup1-GFP were treated with 20 µg/ml methionine for 15 min and washed twice with 400 mM NEM. Cells were lysed in Urea cracking buffer [50 mM Tris-HCl (pH 8.0), 1% SDS, 8M Urea, 20 mM NEM, 1x protease inhibitor cocktail (Roche)] and lysed by beating with 0.5 mm YZB zirconia beads (Yasui Kikai) for 1 min. High salt IP buffer with 20 mM NEM and 0.2% Triton X-100 was added to the lysate, and the samples were rotated at 4°C for 10 min. The solubilized lysates were cleared at 500 × *g* for 5 min at 4°C, and the resultant supernatants were subjected to a high-speed centrifugation at 17,400 × *g* for 10 min. The cleared supernatants were incubated with pre-equilibrated GFP-TRAP_A beads (Chromo Tek) and rotated at 4°C for 1 hour. After the beads were washed with SDS wash buffer [50 mM Tris-HCl (pH 8.0), 250 mM NaCl, 1% SDS, 4M Urea, 5% Glycerol], the bound proteins were eluted by incubating the beads in SDS-PAGE sample buffer at room temperature for 5 min.

### Preparation of Yeast Cell Lysate

Cell lysates were prepared as follows: cells were grown to mid-log phase at 26°C. Aliquots of cells were mixed with trichloroacetic acid at a final concentration of 15%, and the mixtures were incubated for 30 min at 4°C. After centrifugation at 17,400 × g for 10 min at 4°C, the cells were washed once with 100% acetone and then were lysed in Urea clacking buffer [50 mM Tris-HCl (pH 7.5), 8M urea, 2% (w/v) SDS, and 1 mM EDTA] by beating with 0.5 mm YZB zirconia beads (Yasui Kikai) for 5 min. Then, 2x sample buffer [150 mM Tris-HCl (pH 6.8), 7M urea, 10% (w/v) SDS, 24 % (w/v) glycerol, and bromophenol blue] were added to the lysate, and the samples were vortex for 5 min. After centrifugation at 10,000 × g for 1 min at room temperature, supernatants were analyzed by SDS-PAGE and immunoblotting using anti-GFP and anti-G6PDH.

### Quantitative Analysis of Cargo Sorting (Mup1 and Can1-FKBP)

Cell lysates expressed GFP fused cargoes were prepared as described, and then analyzed by immunoblotting. Intensities of the band of processed GFP (GFP’) at 90 min stimulation were measured. Processed GFP in WT was set to 100%. The data and error bars were obtained from three individual experiments.

### Quantitative analysis of GFP-CPS localization

The GFP-CPS localization was classified into two categories: vacuole lumen and vacuole membrane localization. Cells having both vacuole lumen and vacuole membrane localization were classified in the vacuole membrane localization. For each experiment, at least 50 cells were classified. The data and error bars were obtained from three independent experiments.

### Quantitative analysis of Vps10-GFP and Kex2-GFP localization

Vps10-GFP and Kex2-GFP localization was classified into two categories: punctate structures and vacuole membrane localization. Cells having both punctate structures and vacuole membrane localization was classified in the vacuole membrane localization category. For each experiment, at least 30 cells were classified, and the data from three independent experiments were used for the statistical analysis.

### Quantitative analysis of Vps13-GFP localization

Cells having Vps13-GFP punctate structure were quantified. For each experiment, at least 30 cells were classified. The data and error bars were obtained from three independent experiments.

### Quantitative analysis of Vps4-GFP localization

Vps4-GFP puncta colocalizing with mCherry-Vps21 (endosome) was quantified. For each experiment, at least 50 puncta were classified. The data and error bars were obtained from three independent experiments.

### Quantitative analysis of Mup1-BFP localization

Mup1-BFP puncta colocalizing with Vps13-GFP (Vps13 positive puncta) and colocalizing with both Vps13-GFP and DsRed-HDEL were quantified. For each experiment, at least 50 puncta were classified. The data and error bars were obtained from three independent experiments.

### Quantitative analysis of pHluorin fluorescence of Mup1-pHluorin-mCherry

The pHluorin fluorescence of cells expressing Mup1-pHluorin-mCherry was quantified using flow cytometry. For each experiment, 100,000 cells were quantified. The data and error bars were obtained from three independent experiments.

## FIGURE LEGENDS FOR FIGURE SUPPLEMENT

**Figure S1.**
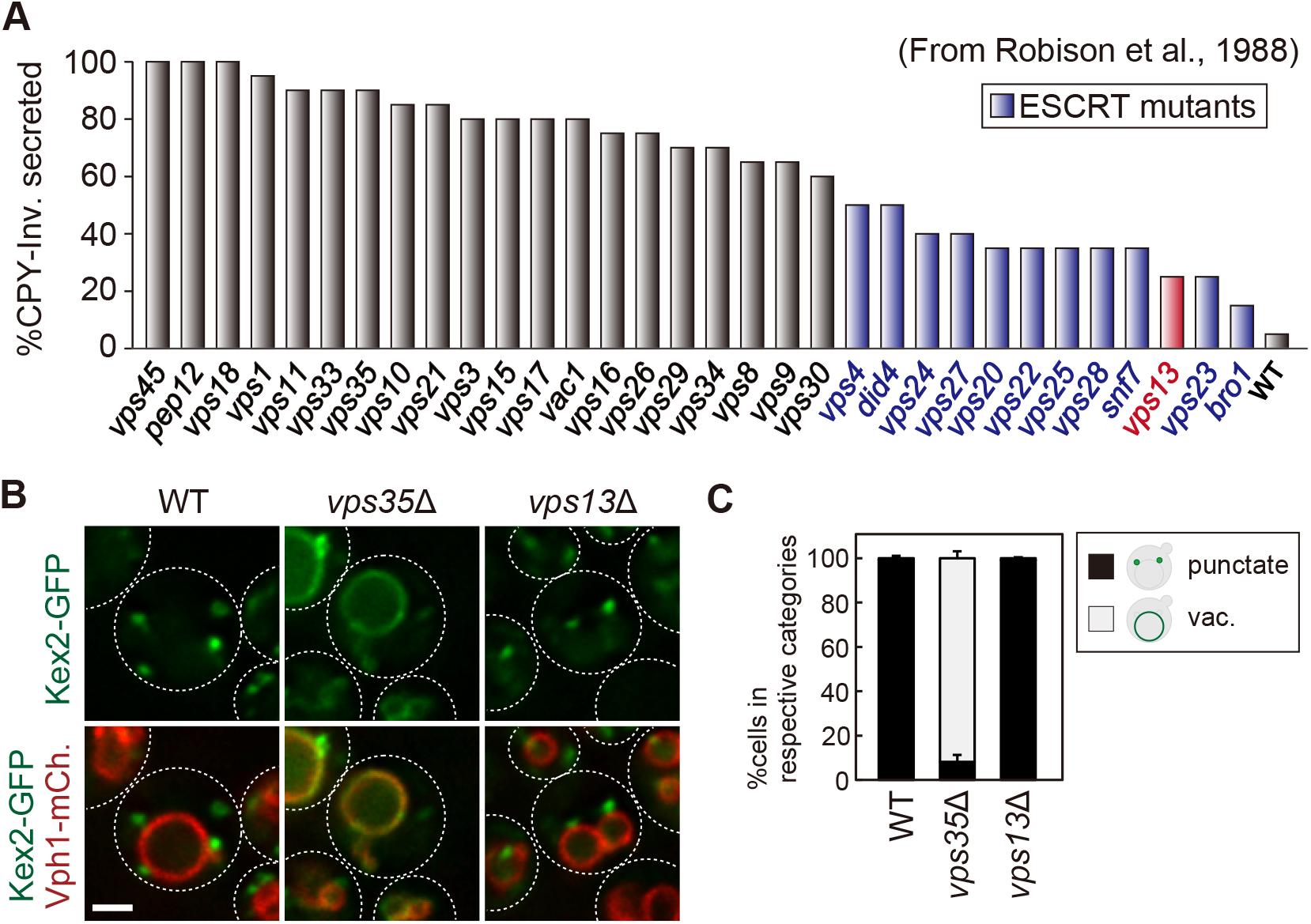
Endosomal sorting pathways in *vps13*Δ cells. (A) CPY missorting index in *vps* mutants. The graph represents the percentage of secreted CPY-Invertase as reported by Robison et al., 1998. (B) Kex2-GFP localization in WT, *vps35*Δ (retromer), and *vps13*Δ cells. (C) Quantification of Kex2-GFP localization. Scale bar: 1 µm.

**Figure S2.**
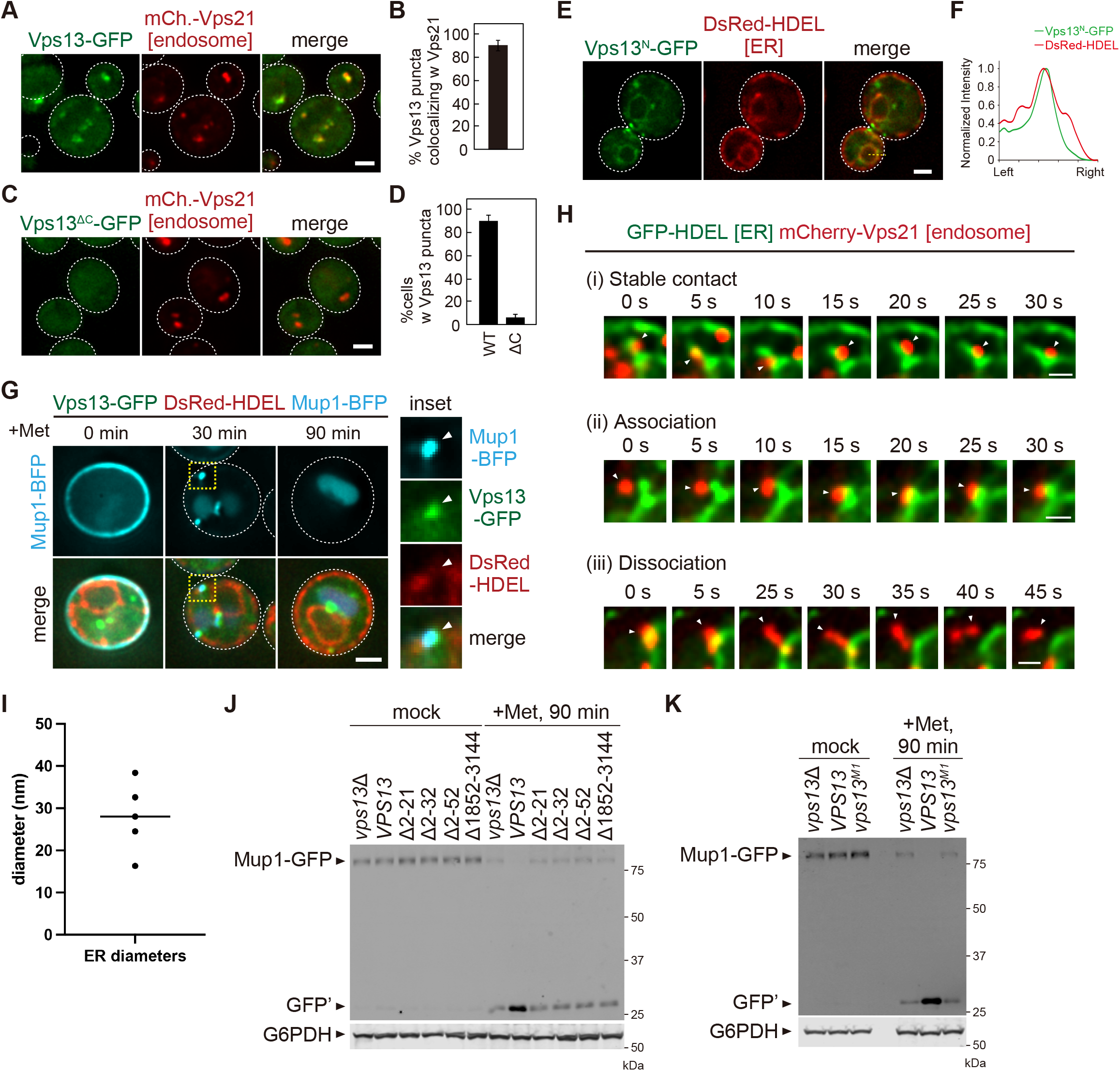
Vps13 localization at the ER-endosome contact site. (A) Localization of Vps13-GFP. Scale bar: 1 µm. (B) Quantification of Vps13-GFP colocalizing with mCherry-Vps21. (C) Localization of Vps13^ΔC^-GFP (residues 1-1851). Scale bar: 1 µm. (D) Quantification of Vps13-GFP puncta localization. (E) The localization of Vps13^N^-GFP (residues 1-39). Scale bar: 1 µm. (F) Line scan analysis for the region highlighted by the yellow dash line in E. (G) Vps13-GFP localization at the ER-endosome contact site. Vps13-GFP, DsRed-HDEL (ER), and Mup1-BFP (endosome) expressing cells were stimulated with methionine. Scale bar: 1 µm. (H) Live cell-imaging analysis of the ER (GFP-HDEL) and endosome (mCherry-Vps21). Scale bar: 500 nm. (I) ER diameters per 100 nm interval starting at the endosome contact along 500 nm length of ER. (J and K) Mup1-GFP processing in *vps13* mutants after methionine stimulation. Cell lysates were analyzed by immunoblotting using anti-GFP and anti-G6PDH antibodies.

**Figure S3.**
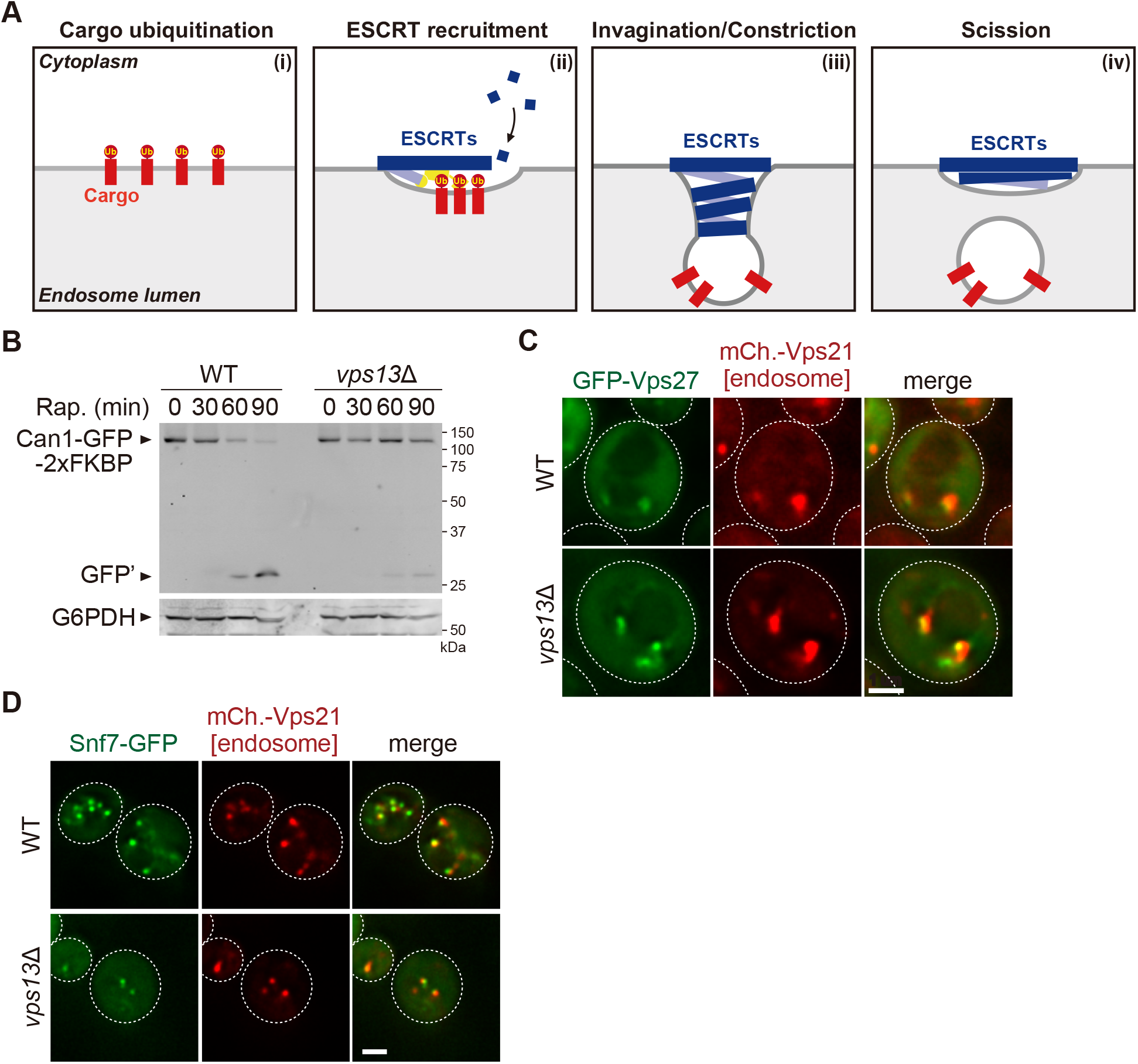
Mup1-GFP sorting in *vps13* mutants. (A) Schematic of ESCRT-mediated sorting at the endosome. (B) Can1-FKBP processing after rapamycin treatments. Cell lysates were analyzed by immunoblotting using anti-GFP and anti-G6PDH antibodies. (C) GFP-Vps27 (ESCRT-0) localization. (D) Snf7-GFP (ESCRT-III) localization. Scale bar: 1 µm.

**Figure S4.**
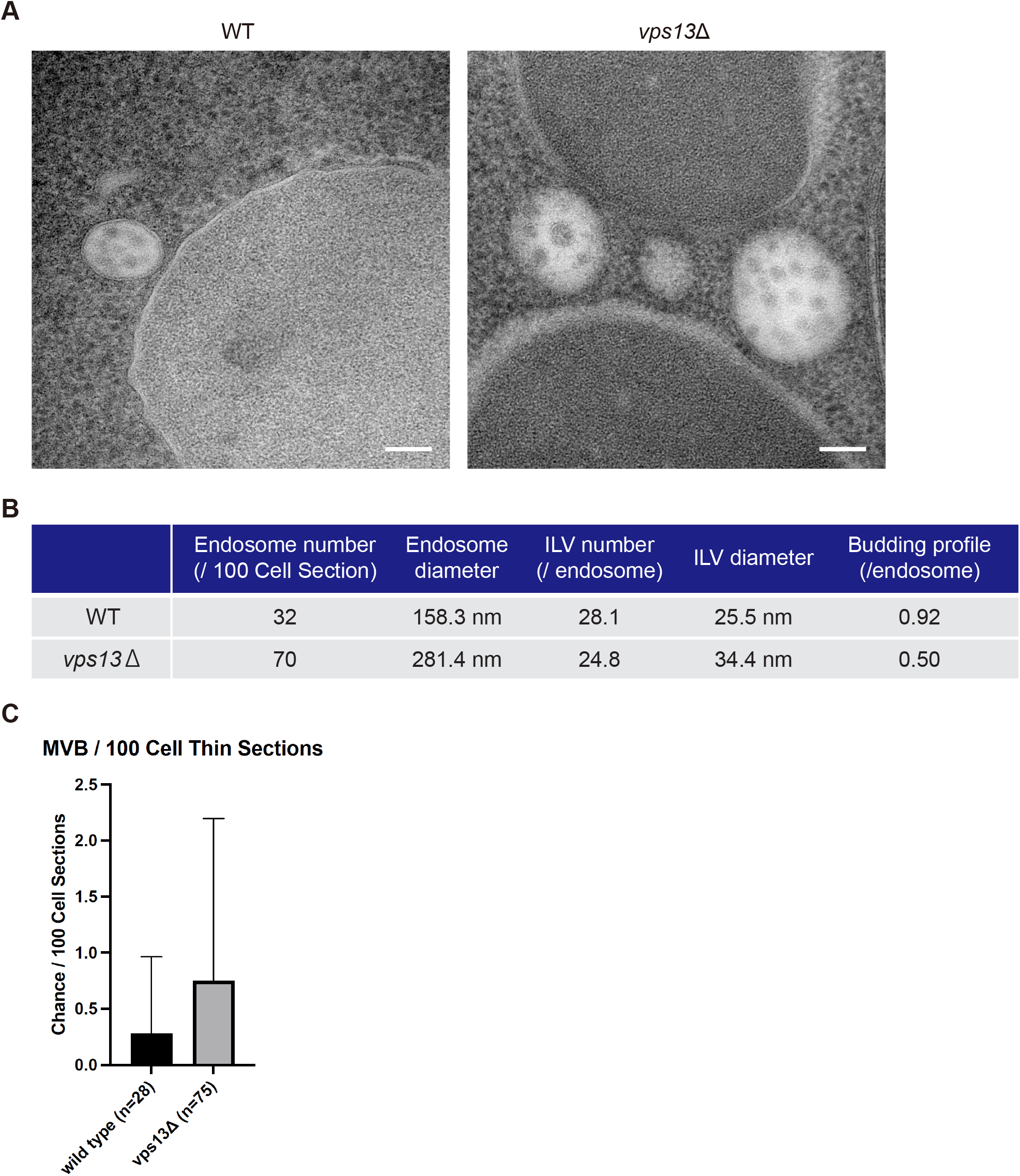
Thin section electron microscopy in *vps13*Δ mutants. (A) Thin section electron miscopy images of WT and *vps13*Δ yeast cells. Scale bars: 100 nm. (B) The value of quantification data used in 4C, 4G, 4H, 4I, and 4J. (C) Quantification of the chance of MVBs (endosomes) in WT and *vps13*Δ cells.

## FIGURE LEGENDS FOR SUPPLEMENTAL MOVIES

**Movie S1. Tomogram of ER-endosome contact sites in *vps4*Δ cells.** ER is shown in green. Ribosomes are indicated as green dots. The endosome stacks are shown in different colors to differentiate individual membranes. Round endosomes are traced in yellow. Larger tubular and cisternal structures are in various shades. Scale bars: 100 nm.

**Movie S2. Tomogram of an endosome in WT cells.** The wild-type endosome contains the intralumenal vesicle (ILV). The limiting endosomal membrane is shown in yellow. The ILVs are in red. Scale bars: 100 nm.

**Movie S3. Tomogram of an endosome in *vps13*Δ cells.** The limiting endosomal membrane is shown in yellow. The ILVs are in red. Scale bars: 100 nm.

**Movie S4. Tomogram of endosomes in *vps13*Δ cells.** The limiting endosomal membrane is shown in yellow. The ILVs are in red. Scale bars: 100 nm.

**Table S1.**
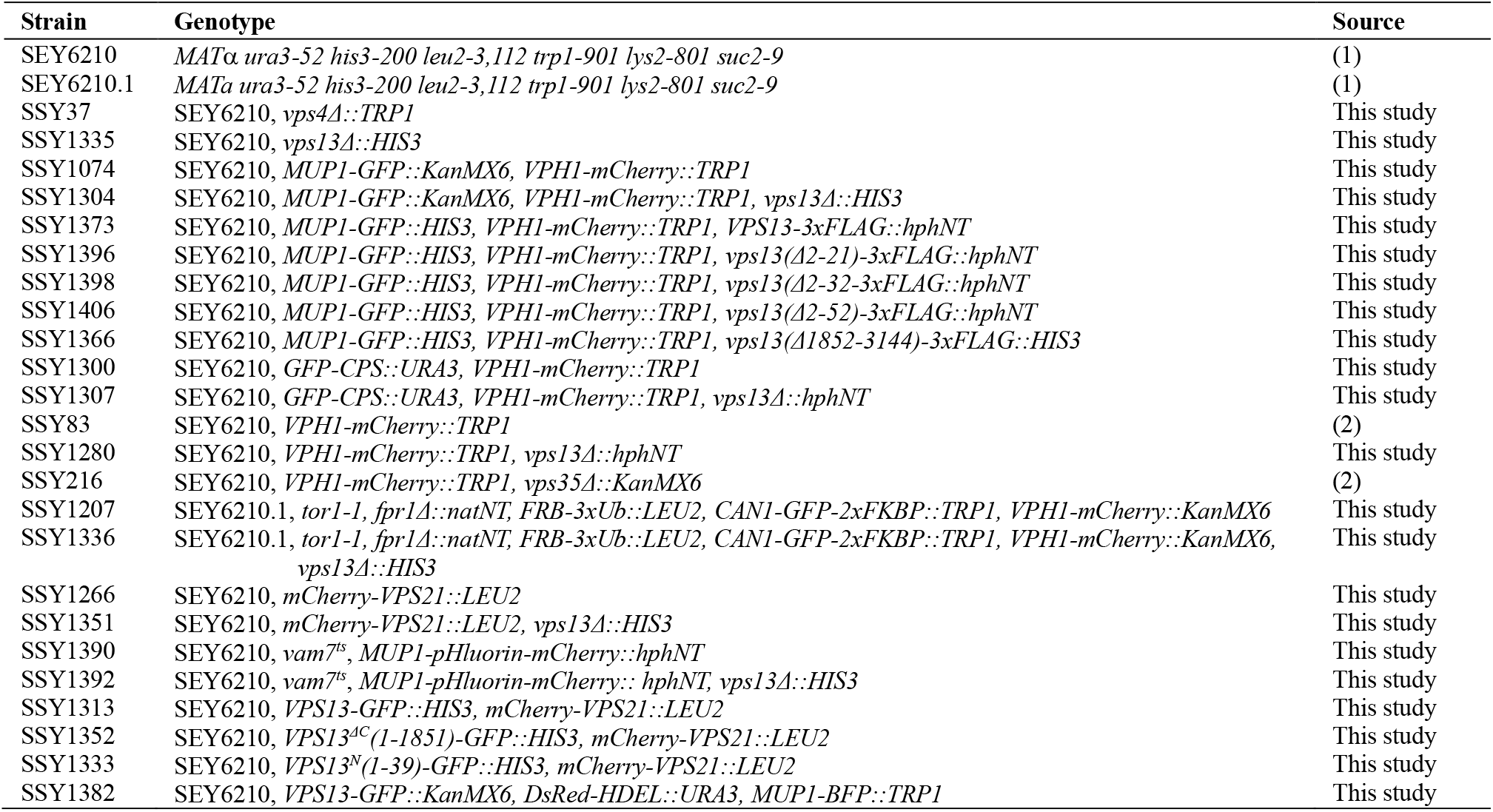
Yeast strains used in this study.

**Table S2.** Plasmids used in this study.

| Name | Genotype | Source |
| --- | --- | --- |
| pRS315 [vec] | *CEN LEU2* | (3) |
| pRS416 [vec] | *CEN URA3* | (3) |
| pRS416-VPS10-GFP | *pRS416-VPS10-GFP* | (4) |
| pRS416-KEX2-GFP | *pRS416-KEX2-GFP* | (2) |
| pRS315-VPS13<sup>^</sup>HA | *pRS315-VPS13pr-VPS13<sup>^</sup>3xHA* | This study |
| pRS315-VPS13(mut1)<sup>^</sup>HA | *pRS315-VPS13pr-VPS13(L64K/I80E/L87E/I162R/L185E/A192E/L217R/V269E/L275D/M293K/L300R)<sup>^</sup>3xHA* | This study |
| pRS416-VPS4-GFP | *pRS416-VPS4pr-VPS4-3xHA-GFP* | (5) |
| pRS426-GFP-VPS27 | *pRS426-VPS27pr-GFP-VPS27* | (6) |
| pRS416-SNF7-GFP | *pRS416-SNF7pr-SNF7-GFP* | (5) |

